# hMT+ Activity Predicts the Effect Of Spatial Attention On Surround Suppression

**DOI:** 10.1101/2024.10.10.617537

**Authors:** Merve Kiniklioglu, Huseyin Boyaci

**Affiliations:** Department of Neuroscience, Bilkent University, Ankara, 06800, Türkiye; Aysel Sabuncu Brain Research Center & National Magnetic Resonance Research Center (UMRAM), Bilkent University, Ankara, 06800, Türkiye; Department of Psychology, Bilkent University, Ankara, 06800, Türkiye; Department of Psychology, Justus-Liebig University Gießen, Gießen, Germany

**Keywords:** spatial attention, middle temporal cortex, motion perception, primary visual cortex, response normalization, surround suppression

## Abstract

Surround suppression refers to the decrease in behavioral sensitivity and neural response to a central stimulus due to the presence of surrounding stimuli. Several aspects of surround suppression in human motion perception have been studied in detail, including its atypicality in some clinical populations. However, how the extent of spatial attention affects the strength of surround suppression has not been systematically studied before. To address this question, we presented human participants with ‘center’ and ‘surround’ drifting gratings and sought to find whether attending only to the center (‘narrow attention’) versus both to the center and surround (‘wide attention’) modulates the suppression strength in motion processing. Using psychophysics and functional magnetic resonance imaging (fMRI), we measured motion direction discrimination thresholds and cortical activity in the primary visual cortex (V1) and middle temporal complex (hMT+). We found increased perceptual thresholds and, thus, stronger surround suppression under the wide attention condition. We also found that the pattern of hMT+ activity was consistent with the behavioral results. Furthermore, a mathematical model that combines spatial attention and divisive normalization was able to explain the pattern in the behavioral and fMRI results. These findings provide a deeper understanding of how attention affects center-surround interactions and suggest possible neural mechanisms with relevance to both basic and clinical vision science.

**Significance Statement:** Our sensitivity to the drift direction of a grating decreases when it is surrounded by an iso-oriented high-contrast surround. Here, we report the first systematic neuroimaging study on how the extent of spatial attention modulates the strength of this ‘surround suppression’. We found that the strength of surround suppression increases as the extent of spatial attention increases, and, pointing to its critical role, hMT+ activity predicts these results. Furthermore, a mathematical model combining spatial interactions and attention could explain the results. In addition to their relevance to basic vision science, these findings have clinical implications because they suggest that the atypical surround suppression observed in some populations could be related to abnormal spatial attention mechanisms.

## Introduction

Sensitivity to a visual stimulus strongly depends on its surround. Notably, discriminating the motion direction of a drifting central grating becomes harder when presented with iso-oriented high contrast surround (Tadin et al., 2003). This perceptual surround suppression phenomenon is frequently attributed to the antagonistic center-surround interactions in the visual cortex, particularly in the middle temporal cortex (Er et al., 2020; Pack et al., 2005; Schallmo et al., 2018; Tadin et al., 2003, 2011; Turkozer et al., 2016). Stimulus characteristics, such as size and contrast, have been shown to influence neural (Pack et al., 2005; Pihlaja et al., 2008; Schallmo et al., 2018; Turkozer et al., 2016; Williams et al., 2003; Zenger-Landolt & Heeger, 2003) and behavioral suppression (Er et al., 2020; Schallmo et al., 2018; Tadin et al., 2011; Turkozer et al., 2016). However, previous research on the effect of attention on surround suppression in human motion processing is relatively limited.

Previous animal studies have shown neural correlates of the attention affecting surround suppression in general, including in motion processing (Flevaris & Murray, 2015a, 2015b; Freeman et al., 2001; Herrmann et al., 2010; Ito & Gilbert, 1999; Maunsell, 2015; Reynolds & Chelazzi, 2004; Sundberg et al., 2009). Human neuroimaging studies, on the other hand, have demonstrated the effect of attention on surround suppression across a variety of features except for motion (Flevaris & Murray, 2015a; Itthipuripat et al., 2014). In a recent behavioral study, we examined the effect of the spatial extent of attention on surround suppression in human motion perception using drifting gratings presented in different size and contrast levels, and found that increasing the spatial extent of attention causes stronger surround suppression (Kiniklioglu & Boyaci, 2022). So far, however, there has been no human neuroimaging study examining how the spatial extent of attention modulates the neural correlates of surround suppression in human motion perception.

To fill this gap, we studied the effect of attention on surround suppression using behavioral and fMRI experiments. For this purpose, we measured motion direction discrimination thresholds and fMRI responses in human V1 and hMT+ using drifting high contrast central gratings presented together with large annular gratings under two attention conditions: a narrow attention condition in which participants were instructed to attend only to the center grating, and a wide attention condition in which they were asked to attend to both the center and surround. To anticipate, consistent with our previous study (Kiniklioglu & Boyaci, 2022), we found stronger perceptual suppression under the wide attention condition compared to the narrow attention condition. Next, we examined whether the strength of neural suppression in V1 and hMT+ reflected this behavioral effect. Finally, we used a mathematical model, namely the normalization model of attention (NMA) (Reynolds & Heeger, 2009) to explain the behavioral and fMRI data and, thus, establish a link between them.

## Methods

### Participants

Ten volunteers (mean age = 25.7 years, eight female) participated in both the behavioral and fMRI experiments of the study. All participants reported normal or corrected-to-normal vision and had no history of neurological or visual disorders. Participants gave their written informed consent before the experiment. The experimental protocols were approved by the Human Ethics Committee of Bilkent University.

### Behavioral Experiment

#### Apparatus

The visual stimuli were presented on a mid-gray background (16.09 cd/m^2^) using a CRT monitor (HP P1230, 22 inches, 1280 x 1024 resolution, 120 Hz refresh rate) in a dark room. Participants viewed the stimuli at a distance of 75 cm, with their heads stabilized by a chin rest. Stimuli were programmed with the Psychophysics Toolbox (Brainard, 1997) on MATLAB 2018b (MathWorks, Natick, MA). A gray-scale look-up table was prepared through direct measurements of the luminance values (SpectroCAL, Cambridge Research Systems Ltd., UK) and used to ensure the presentation of correct luminance values.

#### Stimuli and Design

The stimuli, shown in Figure 1, matched those described previously (Kiniklioglu & Boyaci, 2022). Briefly, we used drifting sinusoidal gratings (frequency: 1 cycle/degree, speed: 4°/s, starting phase randomized) weighted by two-dimensional raised cosine envelopes, the radius of which defined the stimulus size. The stimulus consisted of a center grating surrounded by an annular grating (except for the center-only configuration; see below). The Michelson contrast of both center and surround gratings was 98%.

**Figure 1:**
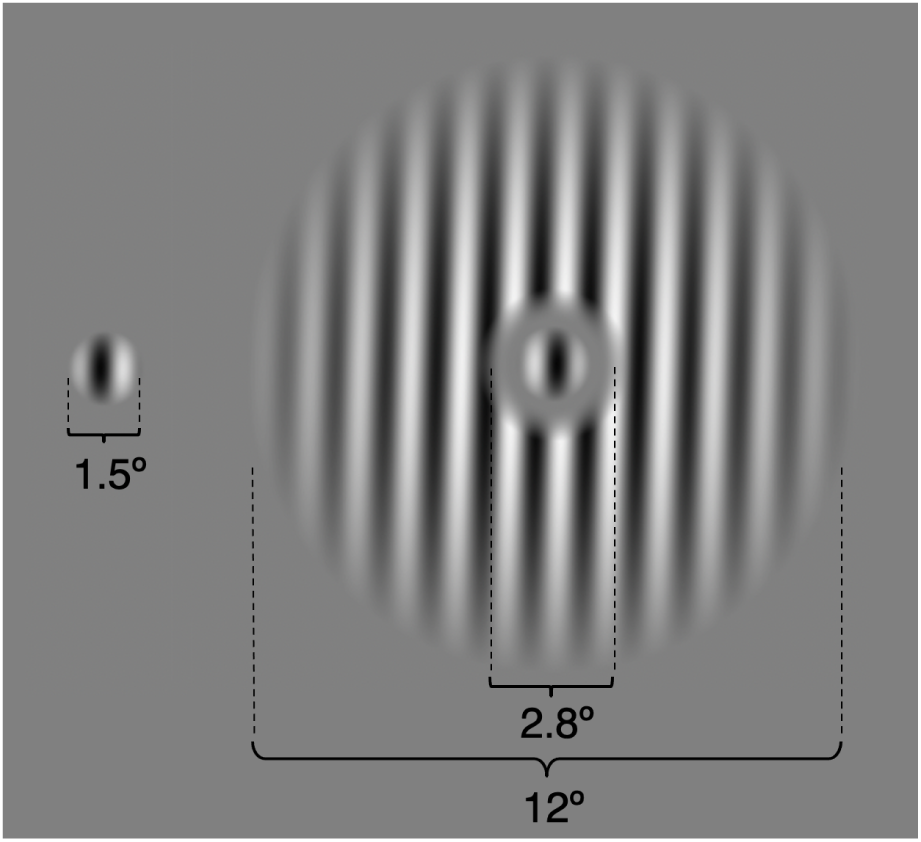
Drifting gratings in center-only and center+surround configurations.

In both the narrow attention and wide attention conditions, the center grating is presented with a surround grating. The center-only trials were used as a baseline to calculate the suppressive effect of the surround on the center. The diameter of the center grating was 1.5°. The inner and the outer diameter of the surround were 2.8° and 12°, respectively. The area between them (1.5° to 2.8°) remained unstimulated to separate the center grating from the surround grating. The center and surround gratings drifted either in the same or opposite direction. For each trial, their directions were determined pseudorandomly such that, in half of the trials, they drifted in the same direction.

Narrow and wide attention conditions were tested in separate blocks, whereas the two direction conditions (same- and opposite-direction) were tested in the same block in randomized order. Center-only trials were tested in the narrow attention blocks. The order of the blocks was counterbalanced for each participant and administered on the same day.

Throughout the experiment, participants viewed foveally presented drifting gratings while maintaining fixation at the center of the stimulus and performed motion direction discrimination tasks via a standard keyboard press. In the narrow attention condition, participants were instructed to attend only to the center grating and report its motion direction. In the wide attention condition, participants were asked to attend both the center and surround gratings. They first reported the drift direction of the center grating, then reported whether the center and surround gratings drifted in the same direction (Figure 2). This second question was solely used to encourage the participants to extend their spatial attention.

**Figure 2:**
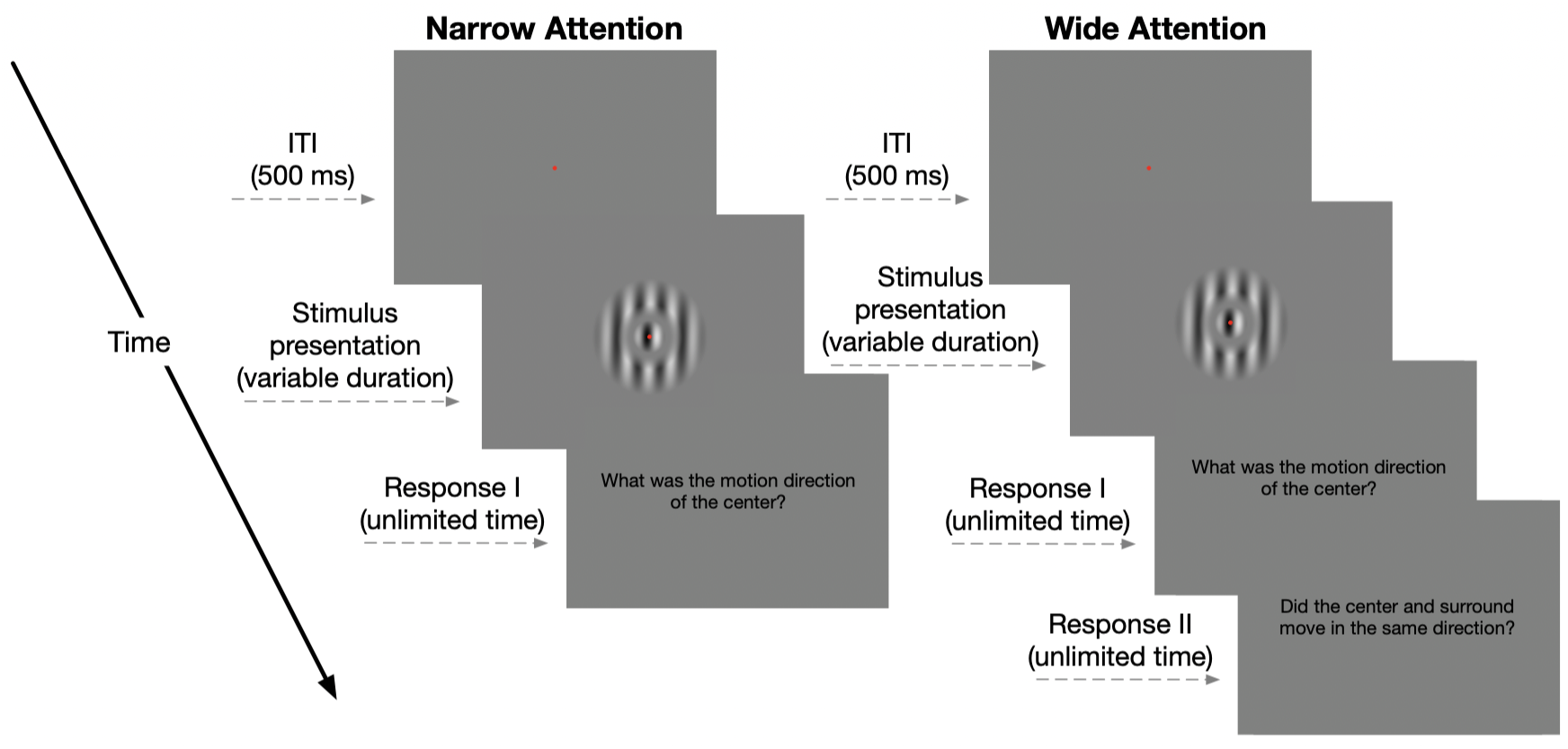
Trial sequence for the narrow and wide attention conditions in the behavioral experiment. Participants were asked to fixate on the center of the display throughout the trial. Each trial began with a fixation point followed by the presentation of the stimulus, the duration of which was adjusted with two interleaved 1-up, 3-down staircases. In the narrow attention condition, participants reported the drift direction of the center grating. In the wide attention condition, participants first reported the drift direction of the center grating and then reported whether the center and surround gratings drifted in the same direction.

In each trial, the duration of the presentation, defined as two standard deviations (SD) of a temporal Gaussian envelope (Borghuis et al., 2019; Tadin et al., 2003), was adjusted with two interleaved 1-up 3-down adaptive staircases, based on the participant’s responses in previous trials. There were two independently progressing staircases for each condition. One staircase started from a very short duration (33 ms), which made the task relatively harder, and the other started from a long duration (158 ms), which made the task relatively easier. There were 100 trials in each staircase. Each participant completed 200 trials for center-only, 400 trials for narrow attention, and 400 trials for wide attention conditions. The experimental session took approximately 45 minutes. Brief break periods were given during the experiment.

#### Data Analysis

Duration thresholds (79% success rate) were estimated by fitting a Weibull function to the proportion of correct responses using the Palamedes toolbox (Kingdom & Prins, 2010) in MATLAB 2019a (MathWorks, Natick, MA) for each participant and condition. Next, using the threshold values, a suppression index, SI, was calculated to quantify the strength of the surround suppression

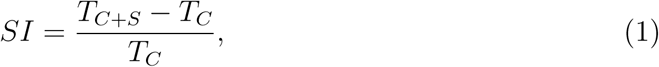

where *T_C_* and *T_C_*_+_*_S_* are the discrimination thresholds for center-only and center+surround configurations, respectively. Higher positive values of SI indicate stronger surround suppression, whereas negative SI values mean surround facilitation. An SI of 0 means no suppression or facilitation. Further statistical tests were performed on the SI values.

First, we compared the SI values to “0” by applying a one-sample, two-tailed Student’s t-test with correction for multiple tests using SPSS Version 25 (SPSS Inc., Chicago, IL). Next, we performed two-way repeated measures ANOVA with factors: attention (narrow and wide attention) and direction (same- and opposite-direction). Then, we conducted two post hoc paired sample t-tests to further explore how the spatial extent of attention affects surround suppression in different direction conditions.

### fMRI Experiment

#### Data Acquisition

MR data were collected on a 3 Tesla Siemens Trio MR scanner (Magnetom Trio, Siemens AG, Erlangen, Germany) with a 32-channel head coil in the National Magnetic Resonance Research Center (UMRAM), Bilkent University. Anatomical data were acquired using a T1-weighted 3-D anatomical sequence (TR: 2600 ms, spatial resolution: 1 mm^3^ isotropic). Blood oxygen level-dependent (BOLD) functional images were acquired with a T2*-weighted echo-planar imaging (EPI) sequence (TR: 2000 ms, TE: 35 ms, spatial resolution: 3x3x3 mm^3^). The stimuli were presented on a 32-inch (1360x768, 60 Hz) MR-compatible LED monitor (T-32, Troyka Med A.S., Ankara, Turkey). The monitor was placed near the rear end of the scanner bore and viewed by the participants from a distance of 156 cm via a mirror attached to the head coil. The stimuli were generated and presented using Matlab and the Psychophysics Toolbox (Brainard, 1997). Participants’ responses were collected via an MR-compatible fiber optic response box (Current Designs). The session started with an anatomical scan, followed by three localizer and six experimental functional runs, and took approximately 1 hour in total.

#### Experimental Runs

The stimuli were drifting sinusoidal gratings as in the behavioral experiment (Figure 1). We used a mixed design in which narrow attention blocks alternated with wide attention blocks. The center-only trials were presented in the narrow attention blocks. The experimental design is depicted in Figure 3. Two conditions of attention (narrow and wide attention) were tested in separate blocks, whereas two conditions of direction (same- and opposite-direction) were tested in the same block in a randomized order. Each run started with a 24s rest, followed by a 168s narrow-attention and a 124s wide-attention task block in a randomized order, interleaved by a 12s rest block, and ended with a 12s rest. Task blocks started with an instruction screen. Within a task block, each condition (three conditions for narrow attention and two conditions for wide attention) was presented six times, and the interval between trials (ITI) was jittered between 4 and 8 seconds. The total duration of a functional run was around 5 minutes. There were six experimental runs in the session.

**Figure 3:**
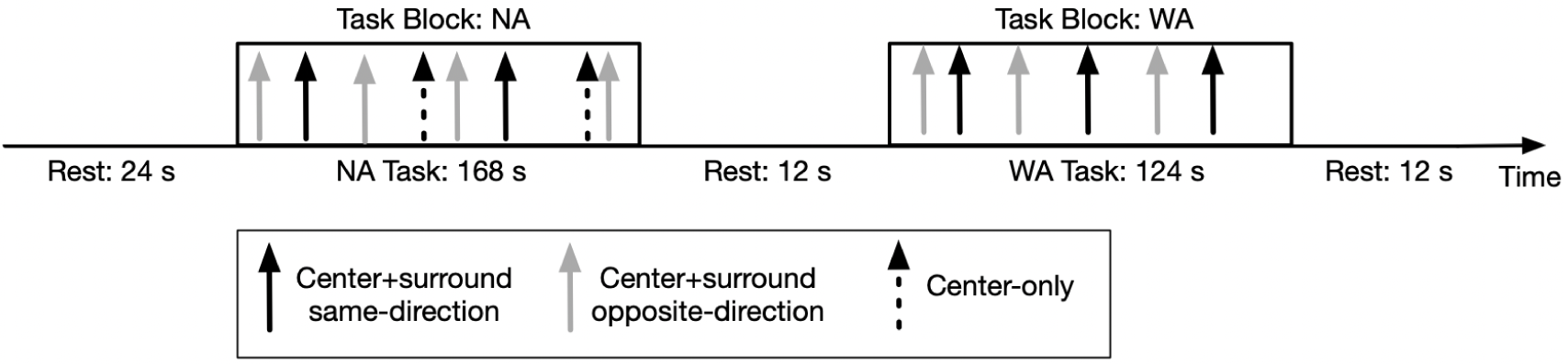
Schematic depiction of the mixed design of the fMRI experiment. In each run, task blocks were alternated with rest blocks. The order of two block types (NA: narrow attention; WA: wide attention) was randomly determined. The narrow attention block consisted of center-only, sameand opposite-direction trials. Wide attention blocks consisted of sameand opposite-direction trials. Within a task block, each condition was repeated six times and the inter-trial intervals were jittered between 4 and 8 seconds.

Stimuli were presented for 150 ms on a mid-gray background. As in the behavioral experiment, the participants viewed foveally presented drifting gratings and performed a task on the perceived drift direction via an MR-compatible response box. In the narrow attention and center-only conditions, participants were asked to attend to the center grating and report its drift direction. In the wide attention condition, the participants were instructed to attend both to the center and surround gratings and report whether the center and surround gratings drifted in the same direction (Figure 4). Unlike the behavioral experiment, we asked only one question in the wide attention condition since asking two consecutive questions could elicit confounding neural activity.

**Figure 4:**
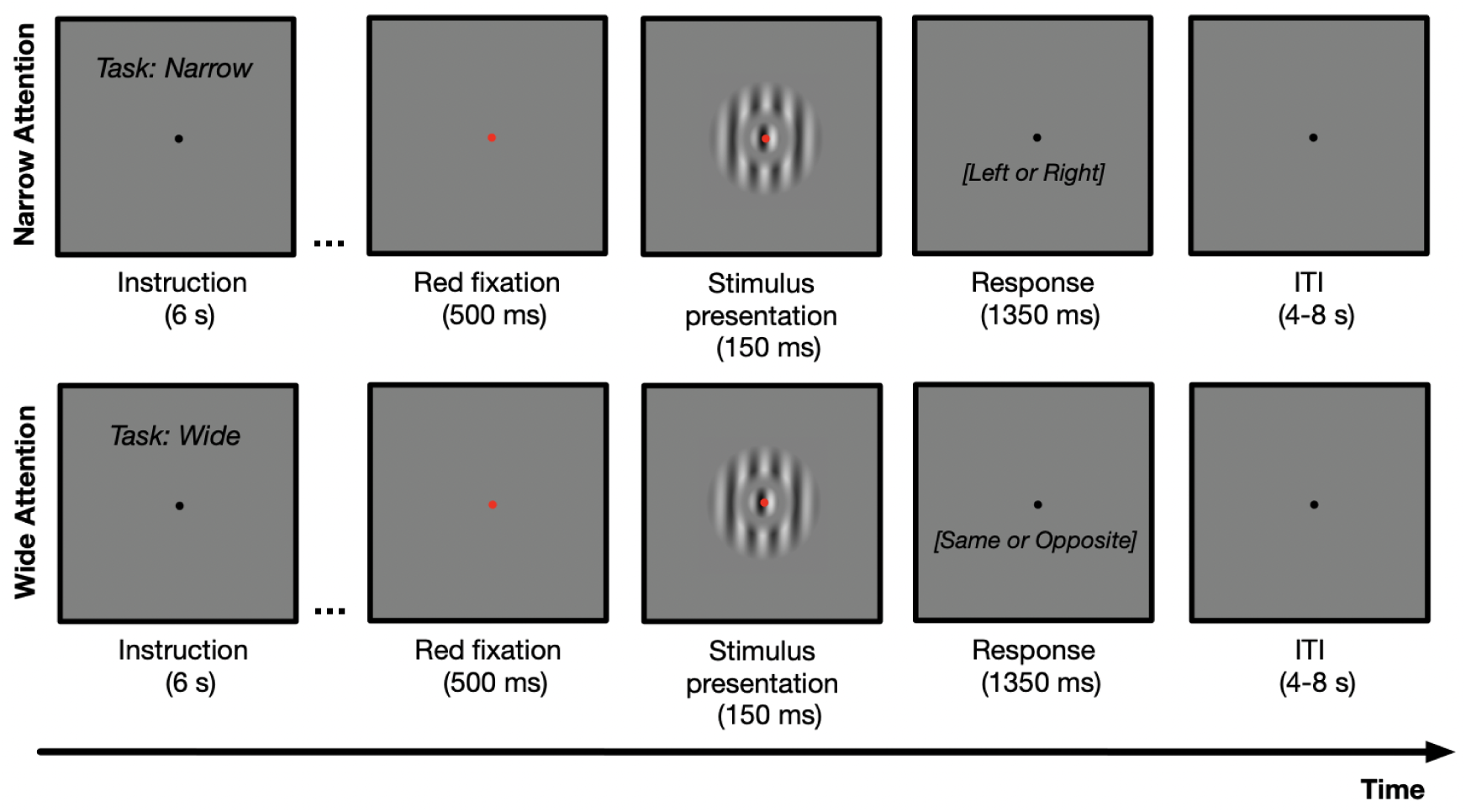
Trial sequence of the narrowand wide-attention blocks in a functional run. Participants were asked to fixate on the center of the display throughout the run. Each block started with instructions about the task. In the narrow attention block, the participants were instructed to attend only to the center grating and report its drift direction. In the wide attention block, the participants were instructed to attend both center and surround gratings and report whether the center and surround gratings drifted in the same or opposite directions. Each trial began with a fixation point, followed by the presentation of the stimulus for 150 ms. In the experiment, possible answers were not written on the screen during the response period.

#### hMT+ and V1 Localizer Runs

To localize participants’ hMT+ areas, we collected fMRI data in a separate run using standard methods in the literature (Huk et al., 2002). Specifically, we used a field of moving and static dots, which consisted of 100 randomly positioned white dots presented foveally within a 12-degree diameter circular aperture on a black background. The dots moved in three different trajectories: cardinal (left-right; up-down), angular (clockwise-counterclockwise), and radial (expanding-contracting). The direction of motion was altered every 2s to prevent adaptation. The run started with a 24s blank period, followed by 12s blocks of a field of static dots alternating with 12s blocks of a field of dynamic dots. There were six dynamic and static blocks in a run. Throughout the entire run, participants were required to maintain central fixation and perform a demanding fixation task in which they were asked to detect changes in the color of the fixation point.

To localize participants’ V1 areas, we collected fMRI data in a separate run. Similar to established retinotopic mapping methods in the literature (Engel et al., 1997; Sereno et al., 1995), we used flickering checkerboard-patterned wedges. However, as we were only interested in the V1/V2 boundary, instead of rotating and expanding wedges, we used alternating horizontal and vertical wedge stimuli (Greenberg et al., 2012; Slotnick & Yantis, 2003). During this localizer run, the presentation of 16s horizontal wedge stimuli blocks alternated with 16s vertical wedge stimuli blocks. This cycle was repeated eight times in a run. Throughout the entire run, participants were required to maintain central fixation and perform a demanding fixation task in which they were asked to detect changes in the color of the fixation point.

#### hMT+ and V1 Sub-ROI Localizer Run

Using an independent localizer run, we identified the set of voxels corresponding to the spatial location and size of the center grating as sub-ROIs within hMT+ and V1. In this localizer run, participants viewed drifting high contrast (98%) center-only and surround-only gratings whose size and location were the same as in the behavioral experiment and the functional runs (Figure 1). The run consisted of 12s active blocks alternated with 12s rest blocks, repeated six times. Center gratings and surround gratings were presented in different active blocks. Throughout the entire run, participants were required to maintain central fixation and perform a demanding fixation task in which they were asked to detect changes in the color of the fixation point.

### Data Analysis

#### Preprocessing

MR data were preprocessed and analyzed using the FMRIB Software Library (FSL) (www.fmrib.ox.ac.uk/fsl) and Freesurfer (Dale et al., 1999; Fischl et al., 1999; Woolrich et al., 2001). High-resolution anatomical images were skull-stripped with BET. Preprocessing steps for functional images included motion correction with MCFLIRT, high-pass temporal filtering (100s), and BET brain extraction. Each participant’s functional images were aligned to her/his own high-resolution anatomical image and registered to the standard Montreal Neurological Institute (MNI) 2-mm brain using FLIRT. Then, the 3D cortical surface was constructed from anatomical images for each participant using FreeSurfer’s *recon-all* command for visualizing statistical maps, anatomical delineation, and identifying ROIs.

#### ROI Construction

For all ROI constructions, a general linear model (GLM) was applied using FSL’s FMRI Expert Analysis Tool (FEAT). Temporal autocorrelations were removed by applying FILM prewhitening (Woolrich et al., 2001).

For the hMT+ ROI, the statistical parametric maps (SPMs) of the dynamic versus static contrast (*α* threshold = 0.05, corrected) were registered to Freesurfer and overlaid on the surface in the native space using the *tksurfer* command. Utilizing the MT label from FreeSurfer’s anatomical delineation for guidance, voxels at the ascending tip of the inferior temporal sulcus and responding more to dynamic compared to static dots were identified as hMT+ and used as a mask for the hMT+ sub-ROI localization.

For the V1 ROI, SPMs of horizontal versus vertical contrast (*α* threshold = 0.05, corrected) were registered to Freesurfer and overlaid on the surface in the native space using the *tksurfer* program. Utilizing the V1 label from FreeSurfer’s anatomical delineation for guidance and voxels that respond stronger to vertical than horizontal wedges, we drew the V1-V2 boundaries. The voxels that fell in or around the calcarine sulcus were identified as V1 and used as a mask for the V1 sub-ROI localization.

For V1 and hMT+ sub-ROIs, we analyzed the independent localizer run data and identified the voxels that respond more strongly to the center compared to the surround grating within V1 and hMT+. The activated regions were identified as V1 and hMT+ sub-ROIs, respectively, and used for further analyses.

#### Analysis of Experimental Runs

Analysis of the experimental functional runs involved extracting GLM beta weights from sub-ROIs using FSL (Jenkinson et al., 2012; Smith et al., 2004). At the first level, we conducted event-related voxelwise analyses, where the predicted fMRI response in each trial was computed assuming a double-gamma hemodynamic response function (HRF). Nuisance regressors for linear motion (derived from MCFLIRT) were also included in the model. For removing the temporal autocorrelations, FILM prewhitening was applied (Woolrich et al., 2001). Contrasts were calculated, focusing on the main effect of each condition. The second level of the functional analysis involved a fixedeffects combination of all six runs for each participant, providing a per-participant average of the first-level contrasts.

Then, to compute the fMRI response within the hMT+ and V1 sub-ROIs, we extracted parameter estimates for each condition using the Featquery tool (FMRIB, Oxford). Specifically, we extracted the beta weights for each trial type across all voxels within each of the predefined ROIs. We treated these extracted beta weights as the fMRI response and performed further statistical analyses on them.

Next, to quantify the changes in fMRI response caused by the inclusion of surround grating and also to draw a link between behavioral and fMRI results, we calculated a Suppression Index (SI) defined as

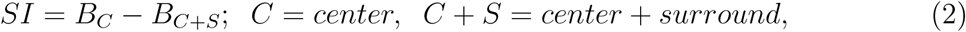

where *B.* is the fMRI response for the given condition. A negative SI means surround facilitation, and a positive SI means surround suppression. An SI of 0 represents no suppression. Further statistical tests were performed on the SI values.

First, we compared the SI values to “0” by applying a one-sample, two-tailed Student’s t-test with correction for multiple tests using SPSS Version 25 (SPSS Inc., Chicago, IL). Next, we performed a two-way ANOVA on the SIs with two factors: attention (narrow attention and wide attention) and direction (same-direction and opposite-direction) for each sub-ROI. Then, we conducted two post hoc paired sample t-tests to further explore how attention affects surround suppression under different direction conditions.

### Model

To explain the behavioral and fMRI results, we used the normalization model of attention (NMA) (Reynolds & Heeger, 2009), which incorporates attention into the computation of neurons’ responses. The model computes population responses as

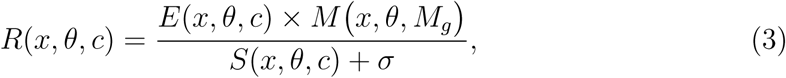

where *x* is spatial position, *θ* is drift direction, *c* is contrast, *E* and *S* are the excitatory and suppressive drives respectively, and *σ* is a semi-saturation constant. The excitatory drive is

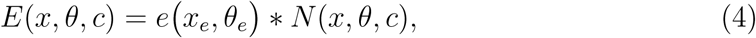

where *e* is a 2-D Gaussian function that determines spatial and direction tuning, *∗* denotes convolution, and *N* is a 2-D Gaussian representing the stimulus. The suppressive drive is

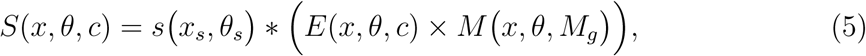

where *s* is a broader 2-D Gaussian tuning function compared to *e*, and *M* is a 2-D Gaussian function whose width is set by the attentional gain factor *M_g_*. Smaller and larger *M_g_*define narrow and wide attention fields, respectively. The operator “*×*” represents element-wise multiplication (Hadamard product). Opposite direction trials are modeled by using surround stimulus direction that is 180° away from the direction of the center (Kiniklioglu & Boyaci, 2022; also see Reynolds & Heeger, 2009). Figure 8 pictorially summarizes the NMA components.

Once the population responses were computed, we used the mean of the maximum responding population as a proxy for the fMRI response, which was calculated by

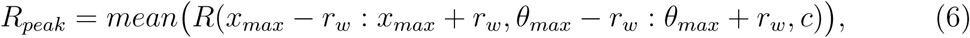

where *x_max_*, and *θ_max_* are the values where *R* attains its maximum value, and *r_w_* defines the width of the averaging area around this peak. Predicted motion discrimination thresholds were then computed as

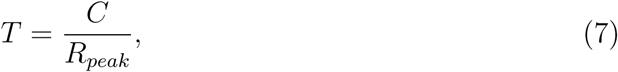

where *C* is a constant linking the behavioral thresholds and neuronal responses.

The model was fit to the data with six free parameters: excitatory spatial pooling width (*x_e_*), suppressive spatial pooling width (*x_s_*), excitatory direction pooling width (*θ_e_*), suppressive direction pooling width (*θ_s_*), attentional gain factor (*M_g_*), and response region width (*r_w_*). Model parameters were estimated using custom functions (Er et al., 2020; Schallmo et al., 2020) and the *fmincon* function of Matlab (version 2022b).

## Results

### Behavioral Results

Figure 5 shows drift direction discrimination thresholds and the suppression indices (*SI*) derived from them. One sample *t*-tests showed that under both attention conditions, the *SI* values were significantly larger than zero in the same direction trials (*p*s *<* 0.001) but not in the opposite direction trials (*p*s *>* 0.05). Moreover, two-way repeated measures ANOVA results revealed a significant main effect of attention (F(1,9)= 5.48, *p <* 0.05, *η*^2^ = 0.38), a significant main effect of direction (F(1,9)= 104.56, *p <* 0.001, *η*^2^ = 0.92), and a significant attention-direction interaction (F(1,9)= 8.17, *p <* 0.05, *η*^2^ = 0.48).

**Figure 5:**
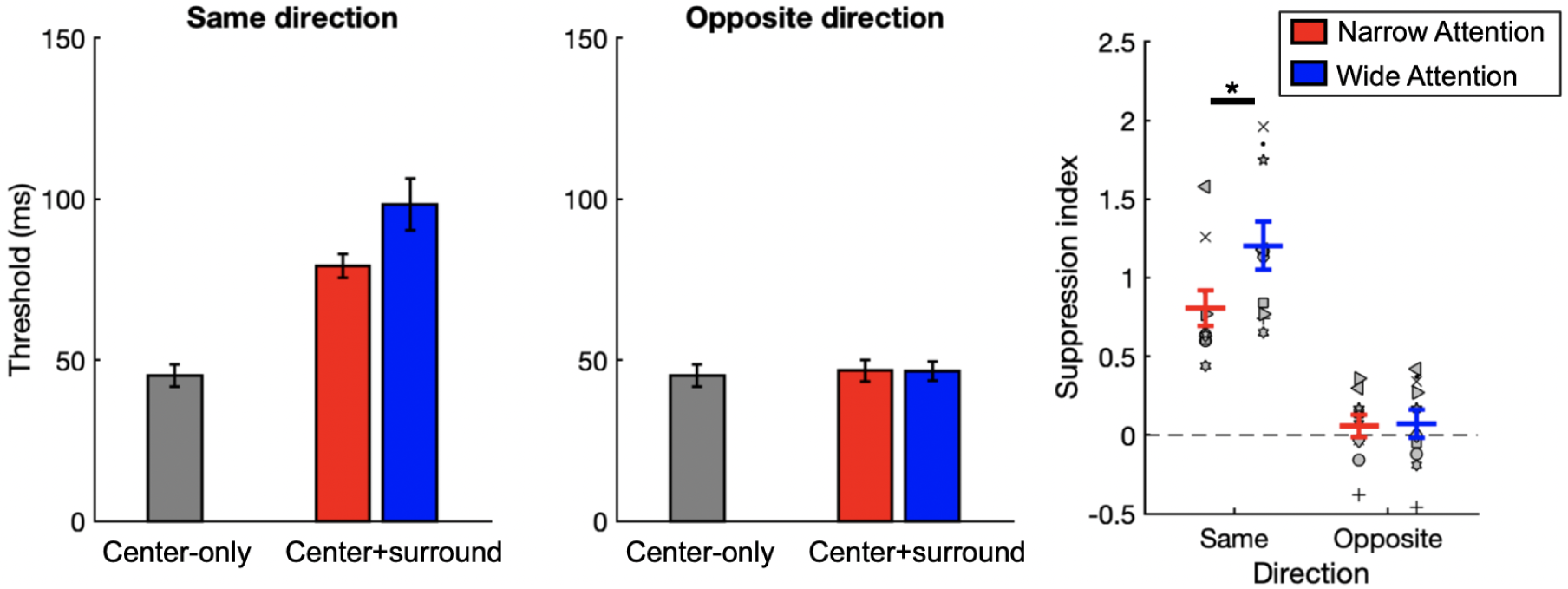
Behavioral results. Left two plots: Drift direction discrimination thresholds averaged across participants. Right plot: Suppression Indices (SI). Higher values of *SI* indicate stronger surround suppression, and negative *SI* values indicate surround facilitation. Symbols represent individual participants, and the red and blue lines show averages for narrow and wide attention conditions, respectively. * indicates significance at *p <* 0.05. Error bars represent SEM.

Post-hoc tests revealed stronger surround suppression under the wide compared to narrow attention condition in the same direction trials, (paired-sample *t*-test: *t* (9) = -2.75, *p* = 0.022; *a_corr_* = 0.025), but not in the opposite direction trials (*t* (9) = -0.24, p > 0.05).

### fMRI Results

Figure 6 shows fMRI responses from hMT+ sub-ROI and suppression indices (*SI*) derived from them. One sample *t*-tests showed that under both attention conditions, the *SI* values were significantly larger than zero in same-direction trials (*p*s *<* 0.0125), not in opposite-direction trials (*p*s *>* 0.05). Moreover, two-way repeated measures ANOVA revealed no main effect of attention (F(1,9)= 1.05, *p >* 0.05, *η*^2^ = 0.10), but there was a significant main effect of direction (F(1,9)= 19.89, *p <* 0.05, *η*^2^ = 0.69), as well as a significant interaction between attention and direction (F(1,9)= 32.81, *p <* 0.001, *η*^2^ = 0.79).

**Figure 6:**
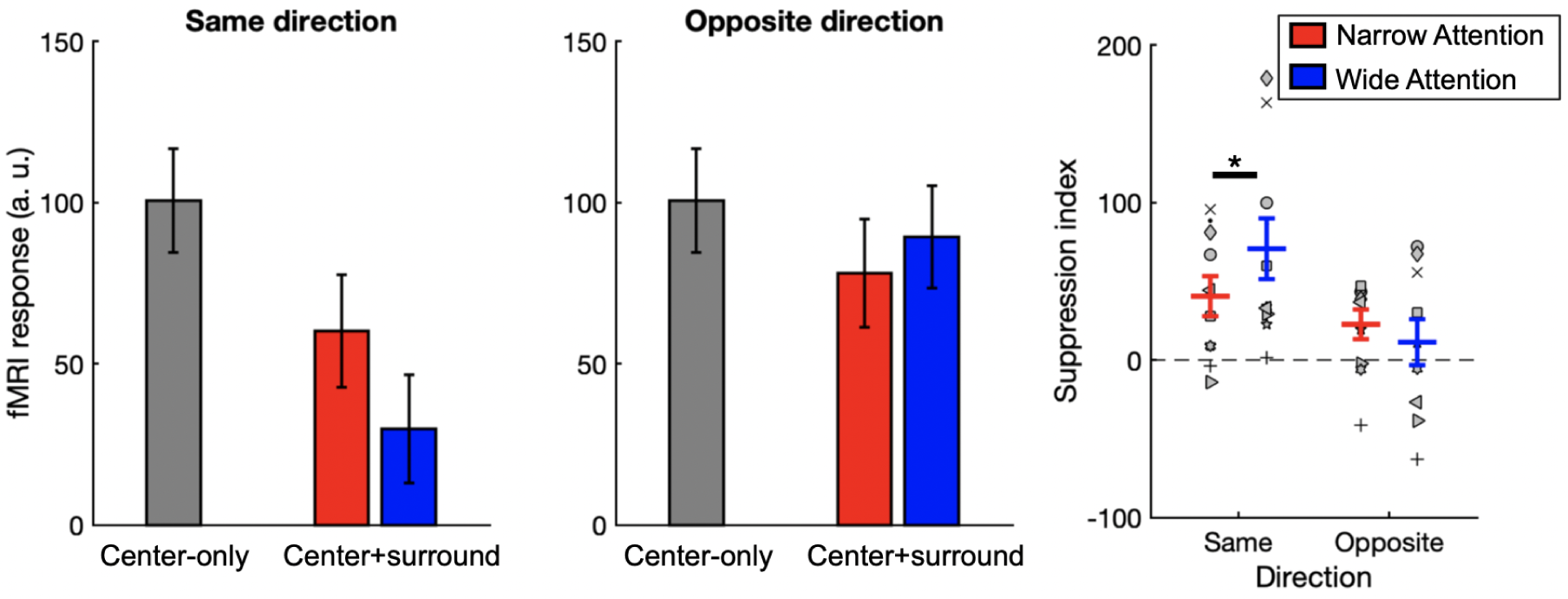
fMRI responses from hMT+ sub-ROI. Left two plots: fMRI responses averaged across participants. Right plot: Suppression Indices (*SI*) derived from fMRI responses. Red and blue lines indicate narrow and wide attention conditions, respectively. * indicates significance at *p <* 0.05. Error bars represent SEM.

Post-hoc paired-sample *t*-test results showed that *SI* values were significantly larger in the wide attention condition compared to the narrow attention condition only in same-direction trials (*t* (9) = 2.90, *p* = 0.018; *a_corr_* = 0.025), but not in opposite direction trials (*t* (9) = -1.18, *p >* 0.05), reflecting the pattern observed in the behavioral experiment.

Figure 7 shows fMRI responses from V1 sub-ROI and suppression indices derived from them. One sample *t*-tests revealed that *SI* values were significantly larger than zero only for wide attention same-direction trials (*p <* 0.01; *α_corr_* = 0.0125). For the opposite-direction trials and same-direction narrow attention trials, *SI* values did not significantly differ from zero (*p*s *>* 0.05). Two-way repeated-measures ANOVA results showed a marginally significant effect of direction (F(1,9)= 4.92, *p <* 0.054, *η*^2^ = 0.35). However, neither the main effect of attention nor the attention-direction interaction reached significance (*p*s *>* 0.05).

**Figure 7:**
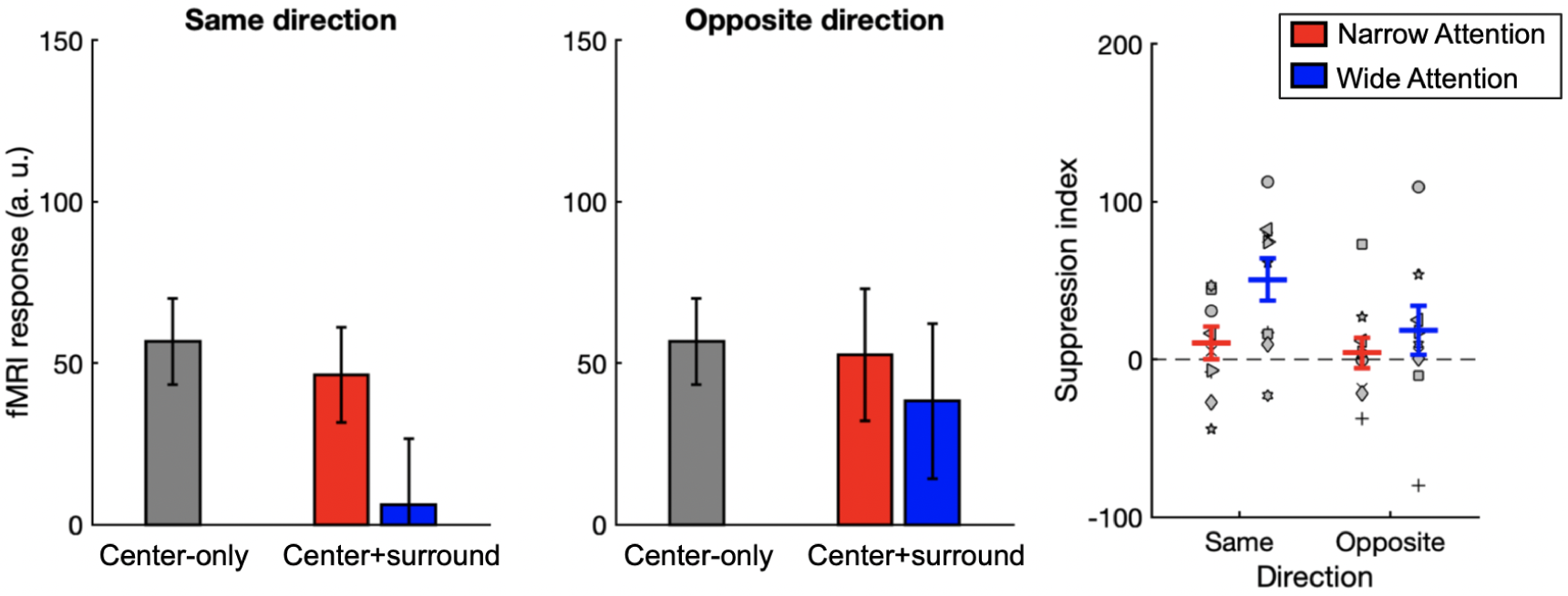
fMRI responses from V1 sub-ROI. Left two plots: fMRI responses averaged across participants. Right plot: Suppression Indices (*SI*) derived from fMRI responses. Red and blue lines indicate narrow and wide attention conditions, respectively. Error bars represent SEM.

**Figure 8:**
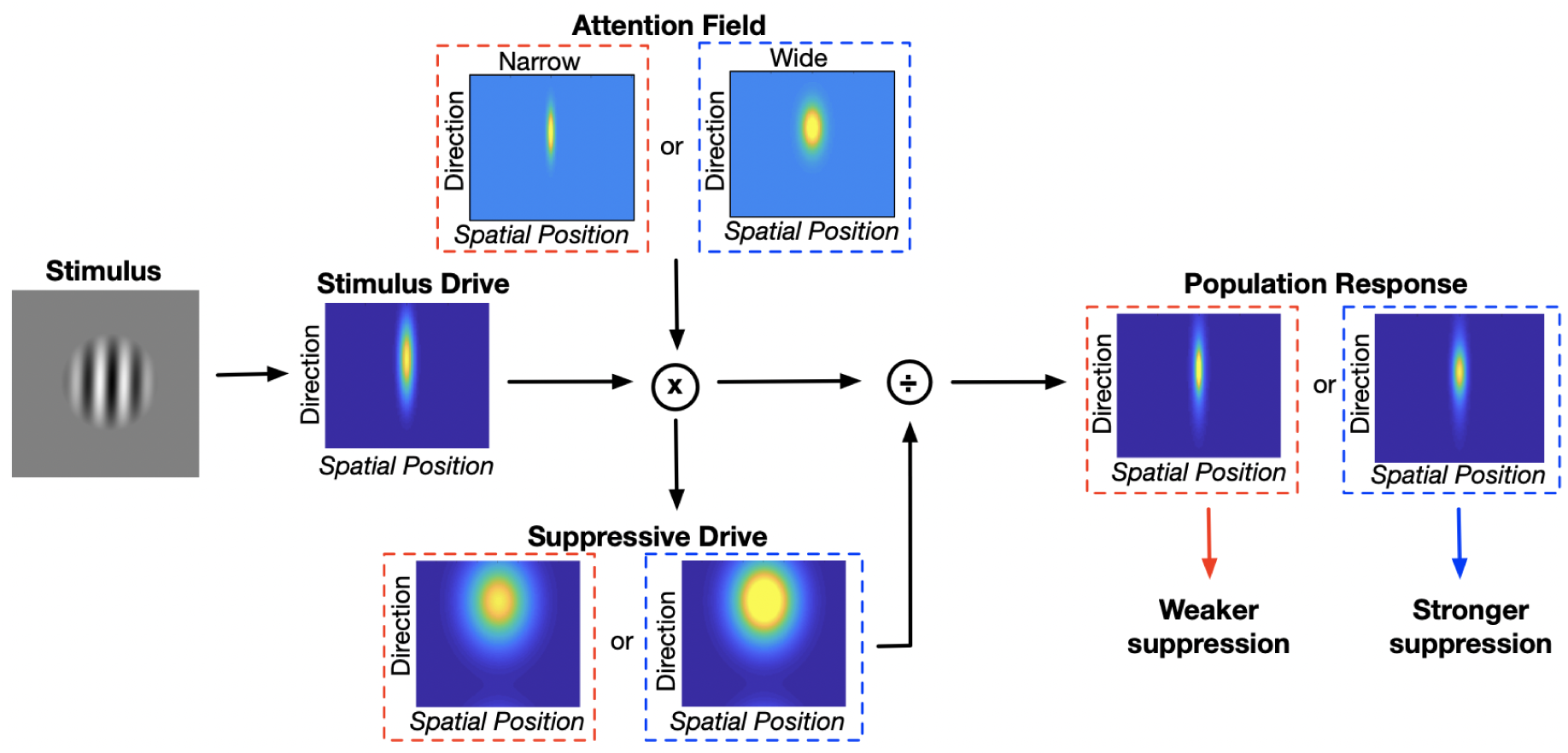
A schematic representation of the normalization model of attention. Stimulus Drive is multiplied by the attention field, then normalized by the Suppressive Drive to determine population response. A narrow attention field leads to a weaker suppression (red dashed squares), and a wider attention field leads to a stronger suppression (blue dashed squares) (Eqs. 3-5).

Post-hoc paired-sample *t*-test results showed that *SI* values were marginally larger in the wide attention condition compared to the narrow attention condition in same-direction trials (*t* (9) = 2.34, *p* = 0.044; *a_corr_* = 0.025), and there was no difference between them in opposite direction trials (*t* (9) = 0.88, *p >* 0.05) Assuming a linear relationship between the fMRI response and neural activity (Boynton et al., 1999) and further assuming that behavioral thresholds decrease monotonically with increased neural activity, our results indicate that hMT+ responses reflect the behavioral effects. However, V1 activity is not consistent with the behavioral data, as we did not find an effect of attention in the same direction trials, and there was no significant suppression in the narrow attention condition. Consequently, our modeling approach in the following section will focus on the hMT+ and behavioral data.

### Model Results

Here, we tested whether we could explain the hMT+ and behavioral responses using the normalization model of attention (NMA) and whether the optimized parameter values are consistent with those derived from previous animal studies. Our implementation of NMA is summarized in Fig 8, and details are described in the Methods section. Briefly, we first compute neural population responses using the divisive normalization model by incorporating a two-dimensional attention field for space and direction. Next, assuming a linear relation between them, we compute the fMRI responses from these neural responses. Finally, we compute the behavioral thresholds from the fMRI responses, assuming an inverse relation between the two.

Table 1 shows the optimized parameter values, and Figure 9 shows the model predictions. The model successfully fits the hMT+ data and qualitatively predicts the pattern of behavioral results. Furthermore, consistent with the task, the optimized value of *M_g_* is smaller for the narrow attention and larger for the wide attention conditions (Table 1). Critically, the optimized model parameters closely match those estimated from the results of previous monkey studies, which are reported in the last column of Table 1. Notably, the values listed in this column represent the average of the parameters derived from several similar studies in the literature (Er et al., 2020; Kiniklioglu & Boyaci, 2022; Schallmo et al., 2018, 2020).

**Figure 9:**
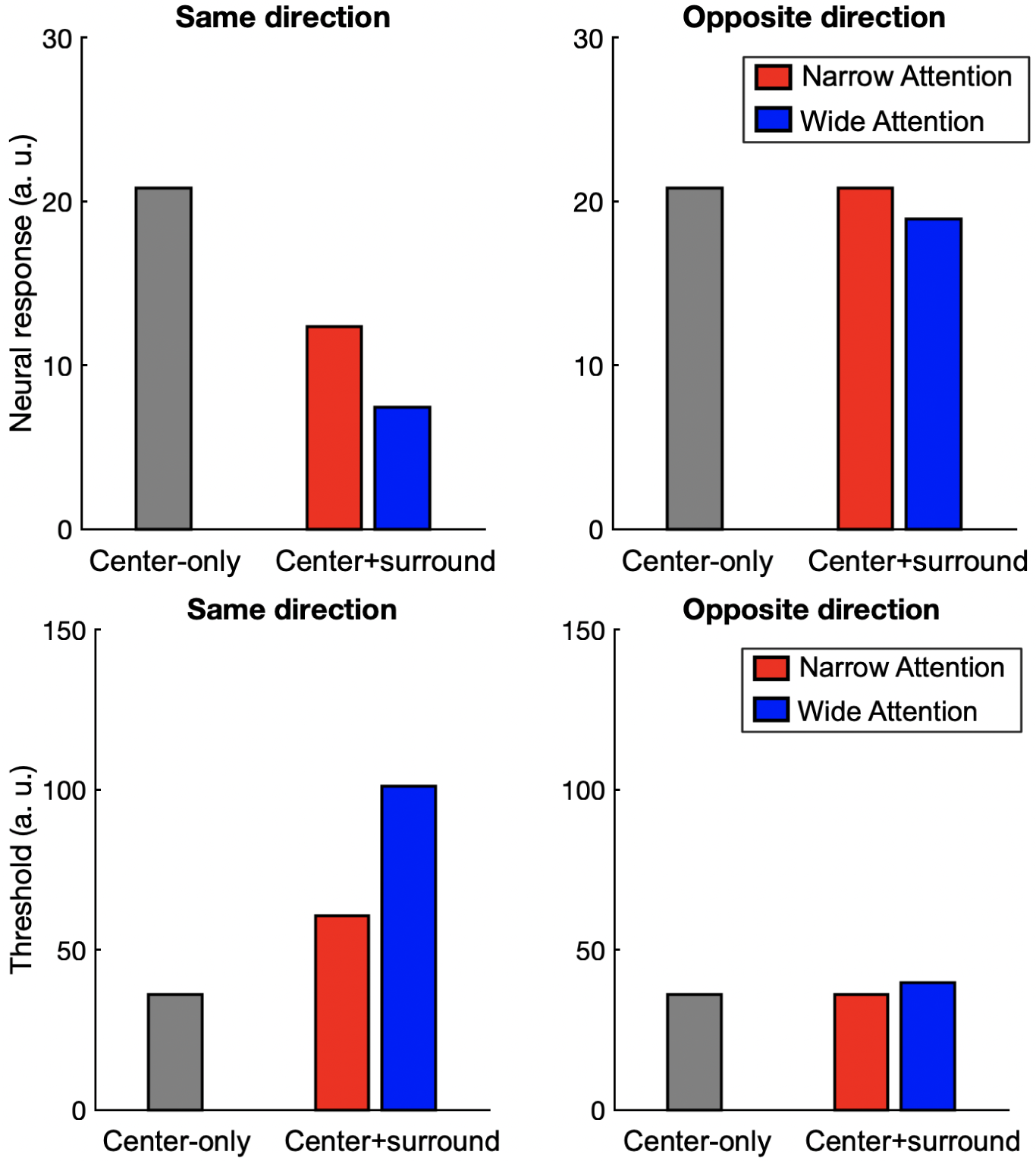
Model results. Top row: Model fit to hMT+ data. Bottom row: Model’s predictions of the behavioral results.

**Table 1:** Normalization model parameters. The last two columns show the optimized values of the parameters in this study and the values estimated from previous monkey neurophysiology studies, respectively.

| Symbol | Description | Optimized | Literature |
| --- | --- | --- | --- |
| $x_e$ | Excitatory spatial pooling width | 3.72 (a.u.) | 4.5 (a.u.) |
| $x_s$ | Suppressive spatial pooling width | 39.86 (a.u.) | 36.3 (a.u.) |
| $\theta_e$ | Excitatory direction pooling width | 24.24° | 23.8° |
| $\theta_s$ | Suppressive direction pooling width | 50.22° | 47.5° |
| $M_g$ | Spatial width of attention field (narrow) | 3.67 (a.u.) | - |
| $M_g$ | Spatial width of attention field (wide) | 25.12 (a.u.) | - |
| $r_w$ | Response region width | 2 (a.u.) | 0 or 1 (a.u.) |

## Discussion

Here, we replicated our previous behavioral findings (Kiniklioglu & Boyaci, 2022) and showed that the strength of perceptual surround suppression increases as the extent of spatial attention increases. Specifically, when the center and surround gratings drift in the same direction, temporal direction discrimination thresholds are larger with wider spatial attention compared to narrower spatial attention. There is, however, no effect when the gratings drift in opposite directions. Our fMRI results showed that hMT+ activity is consistent with this pattern: when the center and surround gratings drift in the same direction, fMRI responses to the center grating are smaller with wider spatial attention compared to narrower spatial attention, and there is no effect when the gratings drift in opposite directions. Finally, performing numerical simulations, we show that a mathematical model, namely the divisive normalization model of attention (Reynolds & Heeger, 2009), can explain the hMT+ and behavioral responses, thus provide a link between behavioral and cortical responses.

To the best of our knowledge, these results provide the first neural evidence for the effect of the extent of spatial attention on surround suppression in human motion processing. The general effect of the spatial extent of attention on neural activity in the visual cortex was previously studied (Itthipuripat et al., 2014; Liu et al., 2022). For example, Itthipuripat et al. (2014), studying steady-state visual evoked potentials (SSVEP), showed that focused attention enhances neural signal-to-noise ratio compared to distributed attention. Similarly, a larger neural response in human V1 is observed when attention is directed to the static center grating compared to surrounding flanker gratings. In motion processing, previous animal studies have frequently reported attentional modulation in MT (Anton-Erxleben et al., 2009; Büchel et al., 1998; Lee & Maunsell, 2009, 2010; Ni et al., 2012; Stoppel et al., 2011; Treue & Maunsell, 1996). Notably, Anton-Erxleben et al. (2009) demonstrated that in macaque MT, the strength of surround suppression increases for the attended location compared to the unattended location by increasing the influence of the attended stimulus on the center neuron’s firing rate. Our findings align well with those earlier results and show that similar neural mechanisms may underly the effect of attention on human motion processing.

Surround suppression is reduced or eliminated when the center and surround move in opposite directions. This has been shown in both neurophysiology studies (Allman et al., 1985; Born & Tootell, 1992; Cavanaugh et al., 2002; Kastner et al., 1995; Lamme, 1995) and behavioral studies (Paffen, Alais, & Verstraten, 2005; Paffen, van der Smagt, et al., 2005). Consistent with those studies, under the opposite-direction condition, we did not observe surround suppression, and we did not find an effect of the extent of spatial attention on it in our behavioral and fMRI results. Note that this also ensured that our results were not due to a task-demand artifact. There are, on the other hand, several human fMRI studies reporting that the opposite direction of surround motion facilitates cortical activity for the center (Moutsiana et al., 2011; Takemura et al., 2012). In those studies, however, authors investigated the effect of motion direction using motion aftereffect (MAE) and induced motion paradigms, not surround suppression, which might explain the difference in the results.

Divise normalization and its variant incorporating attention (normalization model of attention, NMA; Reynolds & Heeger, 2009) have been successfully used to model the correlates of surround suppression in humans using model parameter values derived from monkey studies (Er et al., 2020; Kiniklioglu & Boyaci, 2022; Schallmo et al., 2018, 2020). However, the validity of the model using parameter values obtained directly from human neural data has not been shown before. We found that NMA predicts human fMRI and behavioral data simultaneously, and, importantly, the optimized parameter values closely match those derived from monkey studies. These results demonstrate that divisive normalization and NMA are valid tools to model the correlates of surround suppression in humans and that they can establish a link between the perceptual effects and neural responses.

The pattern of activity in V1 does not completely agree with the behavioral results. This could be related to overall surround suppression mechanisms in motion processing. According to the ’MT-hypothesis’, surround suppression in motion processing originates in hMT+, not V1 (Tadin et al., 2003). Thus, V1 activity may not reflect any effect of attention on surround suppression. Our current results and a number of prior fMRI studies support the MT-hypothesis, showing that hMT+ activity mirrors the behavioral outcomes of surround suppression (e.g., Er et al., 2020; Schallmo et al., 2018; Tadin et al., 2011; Turkozer et al., 2016). Whereas findings about V1 activity are mixed. Some studies found suppression in V1, suggesting that the surround suppression effect could be inherited from V1 by hMT+ (Angelucci et al., 2017; Nurminen et al., 2009, 2013; Zenger-Landolt & Heeger, 2003), several other studies reported both suppression and facilitation depending on presentation and attention conditions (Flevaris & Murray, 2015a; Williams et al., 2003). In the current study, because of several limitations, we cannot completely rule out V1’s potential role. One such limitation is related to the fMRI technique. Our voxel size was 3x3x3 mm^3^, which might be too large to distinguish the V1 sub-ROI activity in response to a small center grating, leading to a reduced signal. Furthermore, because of the foveal confluence, the fMRI response from the V1 sub-ROI might have mixed the activity of several other early visual areas. Thus, although we did not find a good agreement between the behavioral and V1 findings, this may not mean the absence of V1’s role in the perceptual effect.

Weaker surround suppression is observed in multiple clinical conditions, including schizophrenia (Tadin et al., 2006), major depressive disorder (Golomb et al., 2009), and autism spectrum disorder (ASD) (Foss-Feig et al., 2013; Schallmo et al., 2020; Sysoeva et al., 2017). The reason underlying this phenomenon, however, is still not fully understood. Considering that attention mechanisms are affected in many clinical disorders (Clayton et al., 1999; Habermann-Paelecke et al., 2005; Kreither et al., 2017; Luck & Gold, 2008), it is possible that attentional abnormalities lead to atypical visual processing in these patients. This idea is supported by a study that reports weaker surround suppression with an increased level of acetylcholine, which is a neuromodulator thought to be critically involved in attentional processing by leading to more focused voluntary spatial attention (Kosovicheva et al., 2012). Moreover, Schallmo et al. (2020) found weaker suppression in ASD patients, and their modeling work suggested that this could be due to narrower spatial attention, which is in line with our findings. Taken together, these results suggest that the reduced surround suppression found in clinical populations may be caused by abnormal attention mechanisms.

## Author Contributions

**Merve Kınıklıoğlu:** Conceptualization, Methodology, Software, Formal analysis, Investigation, Writing original draft, Writing review & editing, Visualization. **Hüseyin Boyacı:** Conceptualization, Methodology, Writing review & editing, Supervision, Funding acquisition.

## Acknowledgements

This work was funded by a grant from the Turkish National Scientific and Technological Council (TUBITAK 120K956) awarded to HB.

## Data Availability

Data will be made publicly avaliable.

